# Performance, but not size, of hindleg weaponry is sexually dimorphic in the giant mesquite bug (*Thasus neocalifornicus*)

**DOI:** 10.1101/2020.08.03.234385

**Authors:** Zackary A. Graham, Nicole Kaiser, Alexandre V. Palaoro

## Abstract

In many species, males possess specialized weaponry that have evolved to confer a benefit during aggressive interactions. Because male weaponry is typically an exaggerated or extreme version of pre-existing body parts, females often possess reduced or weaponry. Although much research has investigated sexual dimorphism in the sizes of such weapons, other weapon components, such as weapon performance or alternative weapon forms can also explain the evolution of weapon sexual dimorphisms. Here, we investigated the allometry and variation of multiple weapon components of hindleg weaponry in the male and female giant mesquite bugs, *Thasus necalifornicus*. Despite theory predicating greater allocation in male weaponry, we found that females allocated more into the lengths of their hindlegs compared to males. Despite this allocation, males possess relatively wider hindlegs, which likely increase area of muscle mass. Indeed, the squeezing performance of male hindlegs was much greater than that of female hindlegs. Lastly, we also described the allometry and variation in a male weapon component, prominent tibial spines, which likely are used to damage competitors during aggressive interaction. Overall, our findings highlight the intricacies of weapon sexual dimorphism and demonstrate the importance of measuring multiple weapon components and not a single measure.

## INTRODUCTION

Animal weapons are remarkably diverse structures, and have widely variable sizes and shapes within and between species (Emlen, 2008; Wiens & Tuschhoff, 2020). Bearing a large weapon usually increases the fitness of the sex that engages in territorial contests (Husak, Lappin, & Bussche, 2009; Lailvaux & Husak, 2014). As such, weapons are typically under stronger selection in males compared to females, generating sexual dimorphism in weaponry (but see Dalosto, Ayres-peres, Araujo, Santos, & Palaoro, 2019; Stankowich & Caro, 2009). One of the most common forms of weapon sexual dimorphism comes from the disproportional increase in sizes of preexisting structures or appendages. For example, throughout Coleoptera, the males of many species have longer forearms and hindlegs used to grapple with competitors during aggression (Katsuki, Yokoi, Funakoshi, & Oota, 2014; Kojima & Lin, 2017; Rink, Altwegrg, Edwards, Bowie, & Colville, 2019; Wiens & Tuschhoff, 2020). Similar weapon size dimorphisms are common in other arthropods (e.g., crustacean claws, weevil rostrums) and vertebrates (e.g., tortoise gular horns, gorilla forearms), demonstrating the apparent competitive benefit of increased weapon size (Rico-guevara & Hurme, 2018; Vieira & Peixoto, 2013).

Although most examples of weapon sexual dimorphism examples come from differences in size, selection can also favor the diversification of other components of weapons, such as the shape or performance (Palaoro et al. 2020a). For example, in Aeglid anomuran crabs, males have disproportionally longer, and wider claws compared to females’ claws. However, other components, such as strength, mechanical efficiency, and shape do not follow the same expected pattern of sexual dimorphism: males are stronger, but they are not disproportionately stronger than females (Palaoro et al. 2020a). Therefore, sexual dimorphism in weaponry is more complex than just changes in the size of structures. In many cases, there are sex difference in multiple components and supporting traits that might influence how effective a weapon is (Claverie, Chan, & Patek, 2011; Okada, Suzaki, Miyatake, & Okada, 2012; Rico-guevara & Hurme, 2018). Ultimately, the relative importance of a weapon components should determine the degree of sexual dimorphism that is exhibited; with greater degrees of dimorphism between the traits that greatly influence competitive success and less dimorphism between less important traits.

To understand how selection has shaped different weapon components, we can study their allometry and covariation patterns (Palaoro, Peixoto, Benso-Lopes, Boligon, & Santos, 2020). In this way, weapon components that exhibit disproportional growth (steep allometric scaling) might show us how each sex is allocating their resources in each component – and also show us how the components covary with one another. For example, the allometric scaling of weapon size may be similar between males and females, demonstrating that males and females allocate similar resources to their weapon size. However, if the performance capacities (i.e. pinching strength, bite force) of the weapon exhibit steeper scaling in males but not females, this suggests that the performance capacities of males are an important trait in competitive bouts. Thus, measuring the allometry of weapon size alone can be inadequate, because components other than weapon size may have been selected for as well (McCullough, Tobalske, & Emlen, 2014).

A second much neglected aspect of weapon allometry is how components vary within and between each sex. There is ample evidence demonstrating that the expression of weapons are tied to the condition of the individual (i.e., condition dependence), which cause trait hypervariability (Greenway et al., 2019; Johns, Gotoh, McCullough, Emlen, & Lavine, 2014; O’Brien et al., 2018). However, evidence for condition dependence has been shown mainly for weapon size, but not for other weapon components. If we consider that weapon components, such as shape and performance, are traits required to overpower rivals during fights, then any deviation from the optimal shape and performance would decrease the chance to overpower rivals (Arnold, 1983). Therefore, weapon components with high variance should be selected against, which ultimately should decrease variance in these components. For instance, in a aeglid crab species that frequently uses its’ claw to grasp rivals, claw shape has smaller variance than in a species that fights infrequently (Palaoro et al. 2020a). Therefore, by comparing the variability between different weapon components, and between sexes, we can learn about the selection pressures that have acted (and act) on different components of an animal’s weapon. For instance, if the size, but not performance of a weapon is the primary trait that determines the outcome of contests, we expect steep allometric scaling and greater variation in sex that competes more intensely. By contrast, if the squeezing, pinching, or pulling (i.e. performance) capabilities of weapons determine the outcome of contests, we expect little to no sex differences in the allometry of weapon size, but positive allometries in the sex that competes more intensely.

Leaf-footed bugs (Hemiptera: Coreidae) are an ideal system to investigate sexual dimorphism of multiple weapon components. Many male leaf-footed bugs have disproportionally large hindlegs which function as weapons during male-male combat. According to descriptions, male contests are repeated bouts of grappling, in which competitors use their hindlegs to inflict damage to their opponent. Typically, males move around each other until the ventral side of their bodies face each other. When this happens, competitors attempt to grapple and squeeze their tibia into the dorsal surface of their opponent (where the wings are). Interestingly, many leaf-footed bugs species possess enlarged spines on their tibia, which likely increase the potential to injure their opponents and their wings. The hindlegs can also be used during in pre-fight displays in some species, suggesting that in some species hindlegs function as signals (Emberts, St. Mary, Herrington, & Miller, 2018; O’Brien & Boisseau, 2018). Therefore, the sizes and force of the males’ hindlegs likely play an important role in the wielder’s success during competition over mating’s (Emberts et al., 2018; Katsuki et al., 2014; O’Brien & Boisseau, 2018). The hindlegs of female leaf-footed bugs are relatively conserved and used primarily for locomotion, although low intensity aggressive interactions have been observed between females, (Eberhard, 1998). Therefore, understanding the degree of sexual dimorphism in each of the components of the hindlegs might give us further insight into the selective pressures the shape animal weapons.

Here, we investigated sexual dimorphism of multiple weapon components in the giant mesquite bug, *Thasus neocalifornicus*. Although male and female *T. neocalifornius* bodies, forelegs, and midleg are similar, there is significant sexual dimorphism in their hindlegs (Fig 1). Because of the likely role that the disproportional hindlegs of male *T. neocaliforicus* have on male mating success, we investigated differences in the allometric scaling between male and female hindleg morphology and squeezing force. Given the reliance on squeezing rivals and the lack of signaling during typical leaf-footed bug contests, we predicted that performance, rather than size would exhibit steeper allometric scaling compared to females. Furthermore, because squeezing performance might be more important than hindleg size, we predicted that variance in the performance of the hindlegs would be lower than variance in weapon size. Additionally, regarding sexual dimorphism in variance, we expect that males variables should also be more variable regarding all weapons components when compared to females. Lastly, we wanted to characterize the dimorphism in a likely important weapon component — tibial spines. Male, but not female *T. neocalifornicus* possess dimorphic tibial spines; either a single (Fig 1A) or double spine (Fig 1C), which may be used to puncture the wings of an opponent during aggression (Z. Graham, personal observation). If spines are important for force production or competitive success, we expect that tibial spines will also exhibit disproportional scaling and variation.

**Figure 1.**
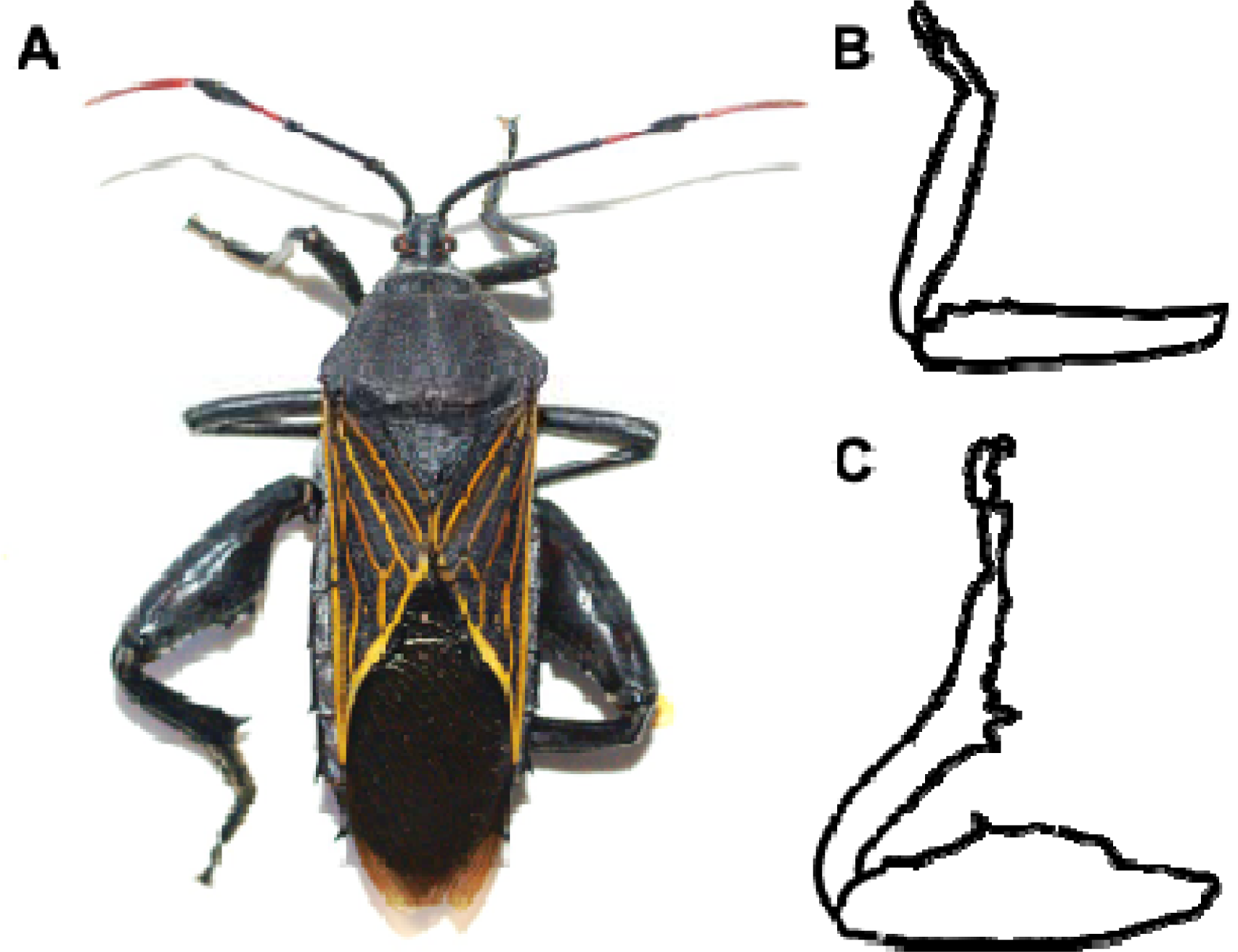
**A)** An adult male giant mesquite bug (*Thasus neocalifornicus*). **B)** Sketch of a typical female and **C)** male hindleg of *T. neocaliforicus*. Males, but not females have exaggerated, widened femurs which can have a single (**A**) or double (**C**) prominent tibial spines.

## METHODS

### Animal collection and care

Adult male and female *T. neocalifornicus* were either hand caught or captured using a net in southern Tucson, Arizona in August and September 2019. In total, we collected 38 males and 21 females that were stored in individual plastic 3deli cups. After collection, all animals were transported to the laboratory. While in the laboratory, *T. neocalifornicus* were kept in incubators with a cycling temperature of 40°C during the day and 25°C at night and coincided with the natural 14:10 light cycle. Each container contained access to water, and a variety of stems and pods from the mesquite trees the bugs were captured (Forbes et al., 2003). All animals were acclimated to laboratory conditions for 48 hours before any procedure to standardize the conditions of the animals.

### Morphology and performance data

We first determined he sex of each *T. neocalifornicus* by identifying males by their larger femur width and pronounced spines on their hind legs (Figure 1). We calculated body size estimated by measuring the protonum width of all individuals with digital calipers (Harvey et al., 2006, Forbes et al., 2003, Greenway et al., 2019, Emberts et al., 2018). Both hindlegs of males and females were then photographed and morphometric data was collected using the ImageJ (ref) software. The morphology measurements collected in ImageJ were femur width and femur length, which were measured from the longest and widest sections of the femur, respectively. (Forbes et al., 2003). In males, we also determined the number of prominent tibial spines to characterize the variation in this weapon component.

To measure the squeezing performance of hindlegs, we collected data on maximal squeeze force that each leg was capable of generating. To do so, we grasped each individual by their protonum and introduced a single hindleg to a custom-built force transducer that is composed of two force plates. The femur of each hindleg was introduced to the bottom force plate of the transducer, triggering a squeezing response from the tibia to pressure to top plate. Each hindleg had its squeezing force measured 3-5 times (Graham, Padilla-Perez, & Angilletta, 2020; Wilson, Angilletta, James, Navas, & Seebacher, 2007). The maximal squeezing force generated for each leg was then used as an overall measurement of squeezing force. Additionally, to ensure our measures were reliable, we measured the squeezing forces of each hindleg on consecutive days.

### Statistical analysis

To analyze the repeatability of each hindlegs squeezing force, we conducted a linear model between maximal force recordings in day 1 and maximal force recordings in day 2. The higher the force of the association between both days, the more repeatable squeezing force is.

Because we predicted that male hindleg weapon size and performance would exhibit steeper allometric scaling than females (greater values of slope, β), we built several linear models using femur length, femur width, and leg squeezing force as our dependent variables. In all those models we used sex as a factor and body size as continuous co-variables. Therefore, we were able to test if males had greater slopes than females in these three weapon components using ANCOVAs. To analyze the allometry in male tibial spines, we first regressed spine length against body size to calculate its allometric coefficient using a linear model. Then, to test if squeezing force is associated to the length of the spine, we regressed maximal leg force (dependent variable) against spine length (independent variable), the spine number (independent variable), and their interaction. We would thus be able to test if spine length and number influenced force output. In these models, body size, femur width, femur length, or force production, and tibial spine length were all log-transformed to our allometry analyses comparable for other allometry studies (Voje, 2015).

Lastly, we wanted to test for differences in the variance of weapon components. To test for sexual dimorphism in variance, we tested whether the residuals of body size and femur length, femur width, and maximal squeezing force was greater in males compared to females using a Bartlett test for equal variances. We conducted all analyses in R (R Core Team, 2019).

## RESULTS

### Repeatability of hindleg squeezing force measurements

The maximal squeezing forces recorded on the first day strongly correlated to the maximal squeezing forces recorded the following day for both male and female *T. neocalifornicus* hindlegs (Fig 2; *r*^2^ = 0.81, *t*_108_ = 1.989, *p* = 0.049).

**Figure 2.**
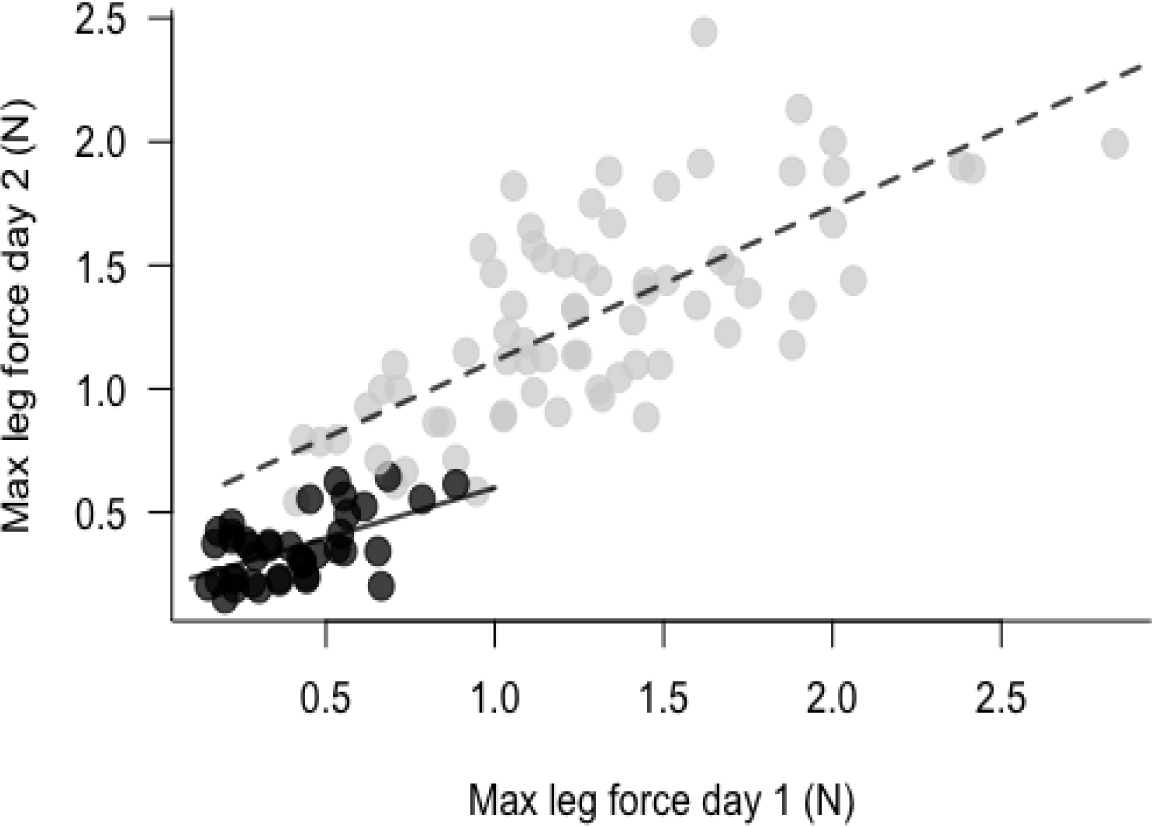
The relationship between the maximal leg force produced on day one and the maximal leg force produced on day two. Black and gray circles represent data from the individual hindlegs of female and male, respectively. The solid and dashed lines represent the best fit line of pinching force repeatability for females and males, respectively. The squeeze forces of both males and females are highly repeatable, which demonstrates that our measurements are reliable measures of force and not measurements of individual variation in squeezing motivation.

### Sexual dimorphism in weapon size and weapon performance

Femur length increased with body size regardless of the sex (Table 1). However, females had greater slopes than males (Fig. 3; females: *β* = 0.95 [0.62; 1.28, 95% CI]; males: *β* = 0.45 [−0.24; 1.15, 95% CI]). We found a similar pattern for femur width, in which femur width increased with body size regardless of the sex (Table 1), and females also had greater slopes than males (Fig. 3; females: *β* = 1.05 [0.58; 1.53, 95% CI]; males: *β* = 0.74 [−0.26; 1.75, 95% CI]). Furthermore, we found that maximal squeezing force production increased as femur length increased regardless of sex (Table 1), although male slope was more than twice as steep in males when compared to females (Fig. 4; females: *β* = 0.95 [−0.69; 2.60, 95% CI]; males: Fig. 4; *β* = 2.54 [−1.33; 6.42, 95% CI]). Similarly, we found that maximal squeezing force increased with femur length regardless of sex (Table 1), but with similar slopes between males and females (Fig. 4; females: *β* = 1.77 [0.28; 3.26, 95% CI]; males: *β* = 1.76 [−1.46; 4.98, 95% CI]).

**Table 1.** Results from the tests of sex differences in weapon morphology and performance scaling performed using ANCOVA.

|  | *d.f.* | F-value | P |
| --- | --- | --- | --- |
| (a) Femur length |  |  |  |
| Body size | 1 | 70.991 | < 0.001 |
| Sex | 1 | 9.125 | 0.003 |
| Body size * sex | 1 | 7.278 | 0.008 |
| (b) Femur width |  |  |  |
| Body size | 1 | 3.99 | 0.048 |
| Sex | 1 | 1742.25 | < 0.001 |
| Body size * sex | 1 | 1.34 | 0.250 |
| (c) Max squeezing force |  |  |  |
| Body size | 1 | 0.199 | 0.656 |
| Sex | 1 | 282.645 | < 0.001 |
| Body size * sex | 1 | 0.002 | 0.962 |

*d.f.*, degrees of freedom.

**Figure 3.**
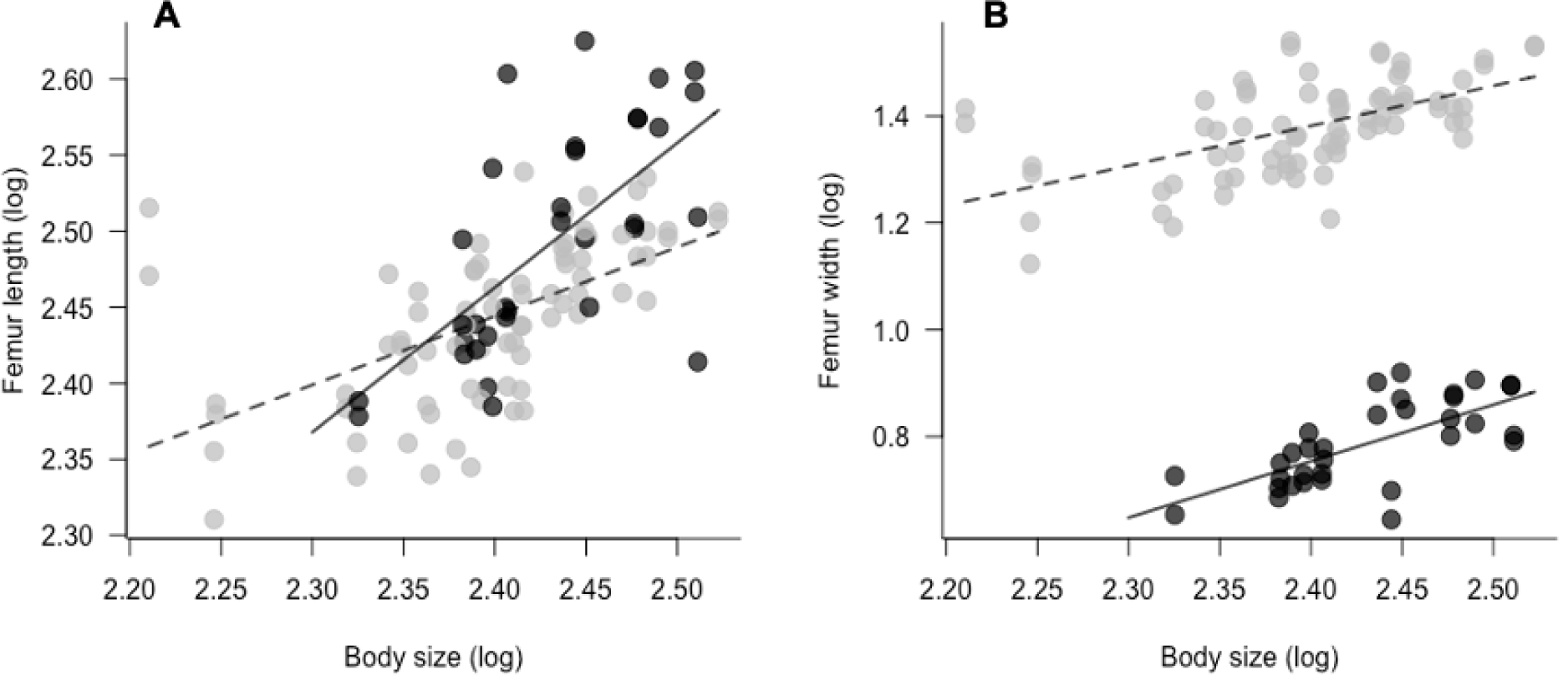
The relationship between body size (protonum width) and hindleg measurements from male and female *Thasus neocalifornicus*. In both figures, black and gray circles represent data from the individual hindlegs of female and male, respectively. The solid and dashed lines represent the best fit line of pinching force repeatability for females and males, respectively. **A)** Male and female body size and femur length overlap substantially, with female having a relatively higher allometric slope. **B)** By contrast, a clear difference arises in allocation to femur width, with males having substantially wider femurs, despite similar allometric slopes to females.

**Figure 4.**
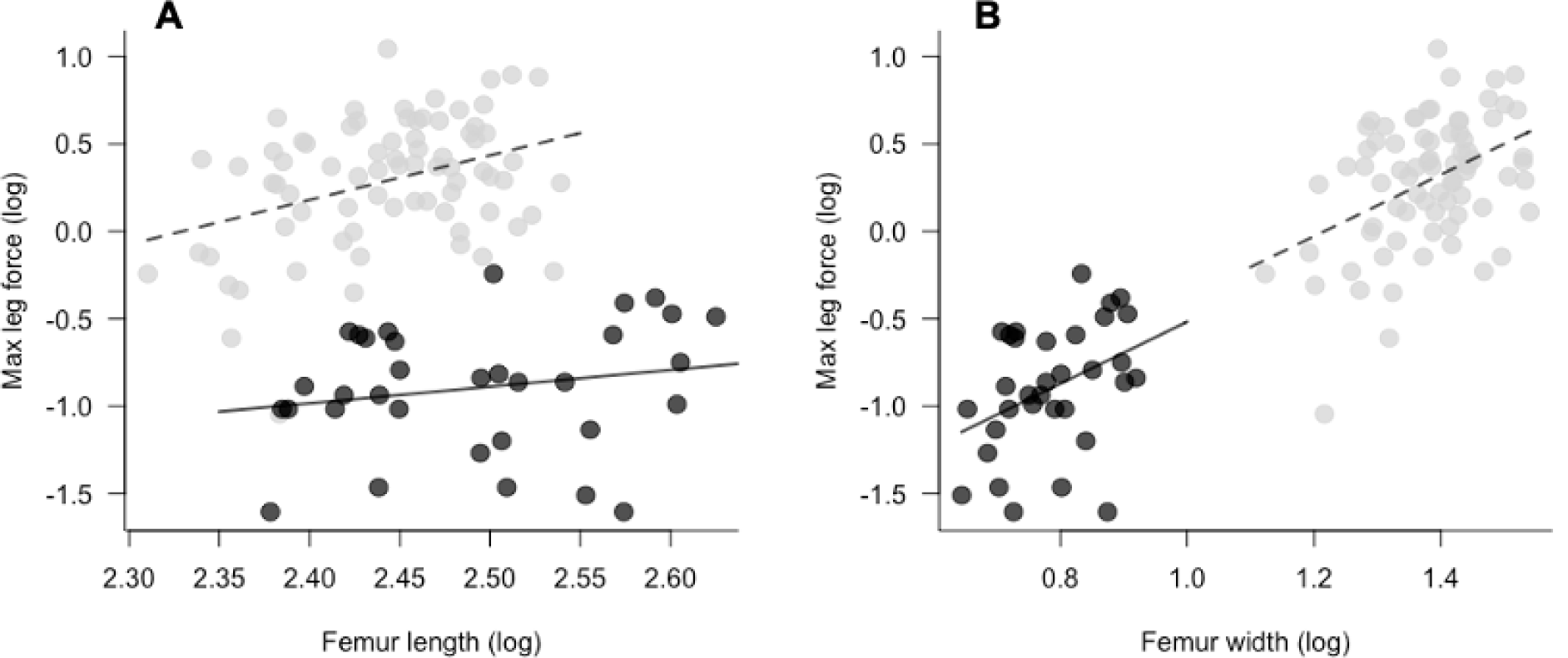
**A)** The relationship between femur length and maximal leg force. **B)** The relationship between femur width and maximal leg force. In both figures, black and gray circles represent data from the individual hindlegs of female and male, respectively. Further, the black and dashed lines represent the best fit line of pinching force repeatability for females and males, respectively. The squeezing forces of male hindlegs are much stronger than the forces obtained from female hindlegs, despite obtaining similar lengths. Variation in force production seems to stem primarily from sexual dimorphism in femur width, which does not overlap between male and female hindlegs.

Regardless of variance of weapon components being similar between the sexes, there were sex differences in the residual variation of body size and the weapon components (Figure 5). The residual variation in femur length was similar between the sexes (Bartlett’s K^2^ = 5.817, p = 0.01; Figure 5). Further, the residuals of body size and femur width in males was more widely distributed than the residuals of females (Bartlett’s K^2^ = 22.231, p < 0.001; Fig. 5). A similar pattern was found for the residuals of body size and maximal leg force: males were more variable than females (Bartlett’s K^2^ = 41.506, p < 0.001; Fig. 5).

**Figure 5.**
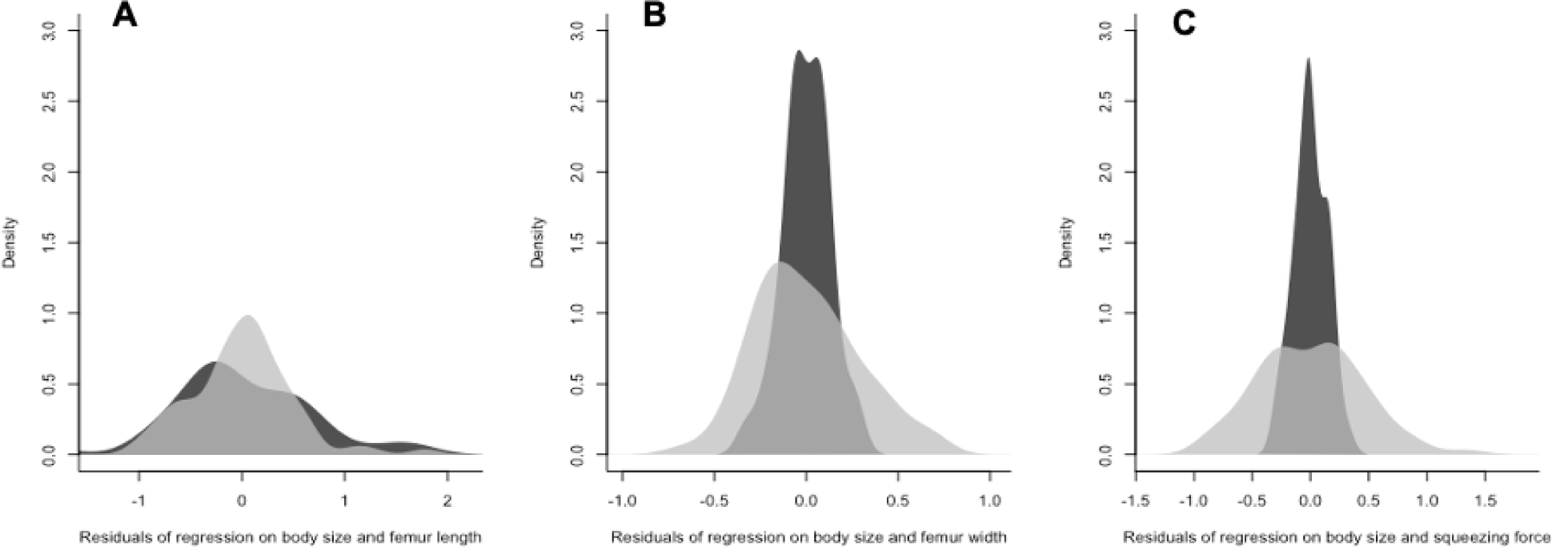
Density plots of the residuals from regressions of male and female **A**) femur length, **B**) femur width, and **C**) squeezing force measurements and body size. In all figures, black areas represent the female data, whereas gray areas represent the male data.

**Figure 6.**
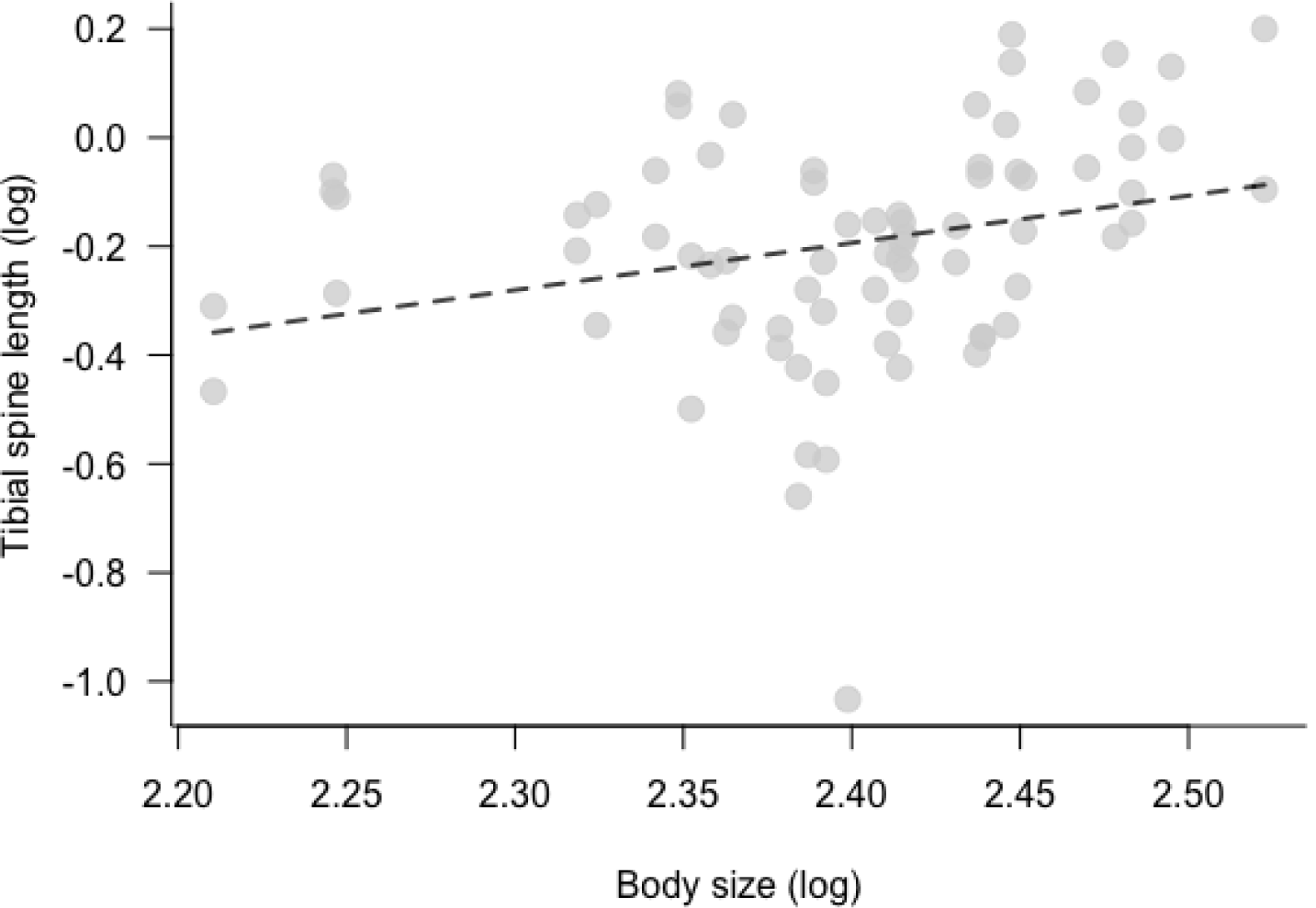
The relationship between body size and tibial spine length in male *Thasus neocalifornicus*. Dashed lines represent the best fit line of body size and tibial spine length. The length of tibial spines increases with body size..

### Male tibial spine allometry

Of the 38 males *T. neocalifornicus* collected, 64 (84.21%) of the male hindlegs had a single tibial spine (Fig 1A) and the remaining 12 (15.79%) hindlegs had a double tibial spine (Fig 1C). Interestingly, of the 12 males with double spine hindlegs, only a single individual possessed double tibial spines on both hindlegs. The remainder of the males either had single tibial spines on both hindlegs, or a mix of one hindleg with a single tibial spine and one hindleg with a double tibial spine.

In males, we found that the tibial spine length increased as body size increased with a shallow slope (β = 0.87 [0.182, 1.556, 95% CI]). When investigating how tibial spine number influenced the squeezing force of the hindleg, we found that there was no relationship between spine number and force production (Fig S3; single tibial spine: β = 0.06 [−0.45, 0.57, 95% CI]; double tibial spine: β = −0.27 [−1.78, 1.23, 95% CI]). However, we did that find tibial spine length increased as force production increased, although the effect was small (β = 0.39[−0.95, 1.72, 95% CI]).

## DISCUSSION

Our results demonstrate the nuances of weapon sexual dimorphism. For example, although theory predicts that males should allocate more into weaponry compared to females, our results support that only some weapon components follow that prediction. Specifically, we found that female hindleg size (both femur length and femur width) exhibited steeper allometric scaling with body size compared to males hindlegs – demonstrating that females allocate more energy into weapon size than males *T. neocalifornicus*. However, despite females exhibiting steeper allometric scaling of the size of their hindlegs, males allocated substantially more into the squeezing performance. Additionally, despite our prediction that male weapon performance would exhibit greater variation when compared to weapon size, we found the opposite for both males and females. Moreover, we also described the allometry and covariation between male hindlegs and the length and number of tibial spines. We discuss these findings and the potential function of tibial spines in the contests of *T. neocalifornicus*.

Although larger weapons are typically associated with contest success (Vieira & Peixoto 2013), increasing weapon size might counterintuitively decrease weapon performance (Levinton & Allen, 2005; O’Brien & Boisseau, 2018). Mechanical lever systems decrease force output as the length of the lever increases (Dennenmoser & Christy, 2013; Swanson, George, Anderson, & Christy, 2013) Thus, because many animal weapons function as lever systems (crustacean claws, leaf-footed bug hindlegs) increasing weapon size can tradeoff with weapon performance. This is known as the ‘paradox of the weakening combatant’, in which large weapons and large muscles are not able to produce higher force output because of the longer levers of larger weapons (Levinton & Allen, 2005). To compensate for this tradeoff, crustaceans and arachnids have tubercles that decrease the length of the lever even in large weapons (Dennenmoser & Christy, 2013; McLain & Pratt, 2011). Although this also seems to be the case here (see below on tibial spines), an alternative strategy would be to decrease weapon size while allocating energy into other weapon components. Here, we found that male *T. californicus* have shorter legs with a lower scaling than females. Additionally, although the scaling for male femur width is lower than females, male intercept (i.e., mean size) is higher than females. Therefore, male femurs are shorter and stouter than females. This morphological pattern implies that male femurs have greater room for muscles than females, which explains the higher performance of males hindlegs.

Outside of the role of *T. neocalifornicus* hindlegs in aggressive interactions, these hindlegs may also be selected for their locomotor abilities. Because the scaling of female hindlegs was near an isometric slope (a slope of 1), this implies that female hindlegs are selected to remain proportional to their body size, enabling efficient and proper locomotion. If the female hindlegs allocated relatively more or less, there may be decreases their locomotor performance. In this way, the weaker allocation in male hindleg length may decrease the locomotor abilities of males. For example, in the frog-legged beetle, *Sagra femorata*, male hindlegs are so large that they are not functional during locomotion (Katsuki et al., 2014; O’Brien, Katsuki, & Emlen, 2017). *S. femorata* males rely purely on their forelegs and midlegs for movement. Perhaps the allometric scaling of male *T. neocalifornicus* is reduced to avoid such the hindlegs becoming nonfunctional for movement, as in *S. femorata*. Therefore, minimal allocating into the size of a weapon may not only increase the mechanical advantage of the weapon but could also influence locomotor performance (Robinson & Gifford, 2019). Future research on how the differential investment of male and female hindleg size influence the locomotor capacity of males and females will shed light on the potential costs and benefits of sexual dimorphic hindleg morphologies.

Despite the size of female hindlegs exhibiting steeper allometric scaling compared to male hindleg size, the maximum squeezing force generated by male legs was much greater than the forces generated by females. The performance capacities of a weapon, and not weapon size, are often important to determine winners in contests (Pinto et al., 2019; De Meyer et al., 2019; Husak, Lappin, Fox, & Lemos-Espinal, 2006). For example, in the male collared lizards, *Crotaphytus antiquus* the outcome of territorial contests are determined by the biting performance of the winner, but not head or body size (Husak et al., 2006). Similar to other leaf-footed bugs, male *T. neocalifornicus* engage in multiple bouts of grappling and attempt to injure their opponents (Z. Graham personal observations). Since females only escalate to low level aggression (which does not involve squeezing rivals, Eberhard, 1998), generating forceful squeezes may not be as important to females. Therefore, it is not surprising that we found much greater allocation in hindleg squeezing force in males compared to females. Indeed, sexual dimorphism in the performance of animal weapons has been studied in multiple taxa, and likely explains the patterns we observed (Goyens, Dirckx, & Aerts, 2015; Goyens, Dirckx, Dierick, Van Hoorebeke, & Aerts, 2014; Husak et al., 2009, 2006; Palaoro et al., 2020; Schenk & Wainwright, 2001; Sneddon, Huntingford, Taylor, & Orr, 2000).

Despite greater allocation by females on weapon size components (i.e., femur length and width), our results on residual variation show that male hindleg force and width are more variable than female hindlegs. By contrast, there was no sex difference in the residual variation in hindleg length or width. Since male *T. neocalifornicus* hindlegs are primarily weapons used to generate forceful squeezes, should selection minimizes the variation in the lengths of male hindlegs. (Levinton & Allen, 2005; O’Brien & Boisseau, 2018). Furthermore, the variation in residual forces from male body size and squeezing force demonstrate that males are much more variable than females. Such increased variation in males is consistent with the hypothesis that squeezing forces and muscle allocation exhibits greater condition dependent in males but not females. Generally, muscle tissue is costly to produce and maintain, and only animals with sufficient resources can invest greatly into greater muscle mass (Somjee, Woods, Duell, & Miller, 2018; Bywater, Seebacher, & Wilson, 2015; Bywater, White, & Wilson, 2014; O’Brien et al., 2019). Overall, by measuring the variation of sexually dimorphic weapon components, we gain a deeper insight into the functional and selection pressures that have acting on such traits.

Interestingly, although the size, shape, and performance capacities of animal weapons are commonly found to be important drivers of weapon evolution, we have also described the polymorphism in an exclusively male weapon component – tibial spines (Fig. 1). Interestingly, the roles of these spines during aggression have not been investigated, but preliminary observations suggest males injure opponents during contests (Z. Graham, personal observation). Sharp, spine-like weapon components are common in many species, such as Gladiator frogs, which possess a spine on their forelegs that are used to injure opponents during contests (Candaten, Possenti, Mainardi, Carvalho, & Palaoro, 2020). Similarly, male *T. neocalifornicus* with double tibial spines may be able to inflict more damage to opponents compared to males with single spines, increasing their chance of winning a contest. Interestingly, the addition of spine-like weaponry is likely to be metabolically cheap way to improve the damage dealing capacities of the weapon. Although the capacity of weapon to deal damage can also be increased through greater allocation into muscle mass, this investment is likely to add substantial metabolic costs (O’Brien et al., 2019). Further, we found that the lengths of tibial spine length increased with body size and force production, suggesting that there may be a relationship between tibial spine length and fighting behavior of *T. neocalifornicus*.

In summary, the variation in weapons, their injury capacities, and how they influence fighting likely promote the diversification of animal weapons (Mccullough, Miller, & Emlen, 2016; Palaoro & Briffa, 2016), but studies of such weapon components influence on contests have received little attention (Palaoro & Briffa 2017, Pinto et al. 2019). Overall, the results of our study highlight the importance of measuring multiple different components of animal weaponry. By only measuring a single weapon variable, such as size, it is difficult to extrapolate on the evolution and complexity of weapon sexual dimorphism.

