## Supplemental Materials for "Performance, but not size, of hindleg weaponry is sexually dimorphic in the giant mesquite bug (*Thasus neocalifornicus*)"

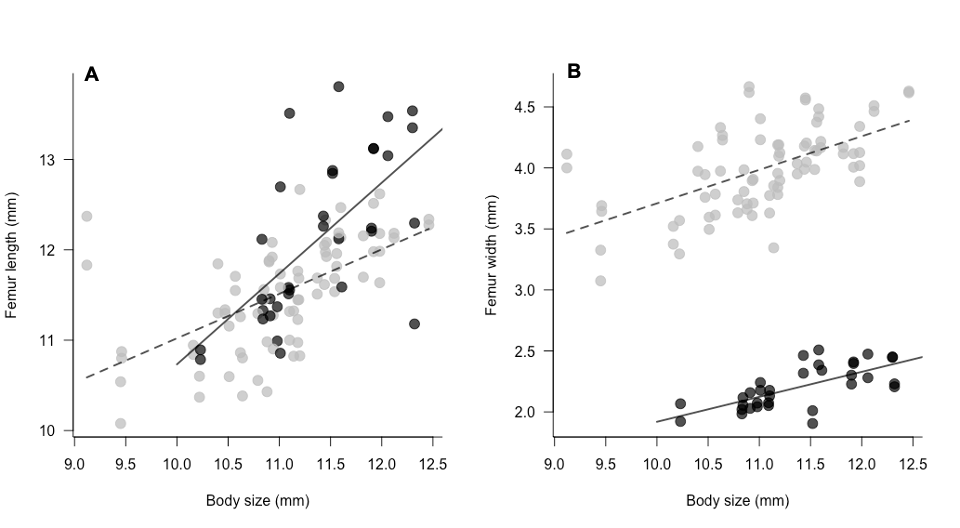

**Figure S1.** The relationship between body size and hindleg measurements femur length and maximal leg force. **A)** In male T**B)** The relationship between femur width and maximal leg force. In both figures, black and gray circles represent data from the individual hindlegs of female and male, respectively. Further, the black and dashed lines represent the best fit line for females and males, respectively.

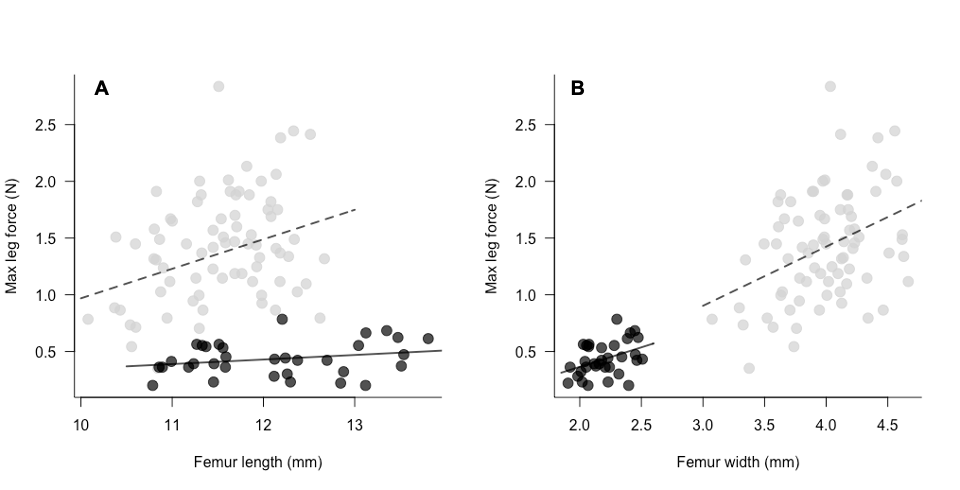

**Figure S2. A)** The relationship between femur length and maximal leg force. **B)** The relationship between femur width and maximal leg force. In both figures, black and gray circles represent data from the individual hindlegs of female and male, respectively. Further, the black and dashed lines represent the best fit line of pinching force repeatability for females and males, respectively. The squeezing forces of male hindlegs are much stronger than the forces obtained from female hindlegs, despite obtaining similar lengths. Variation in force production seems to stem primarily from sexual dimorphism in femur width, which does not overlap between male and female hindlegs.

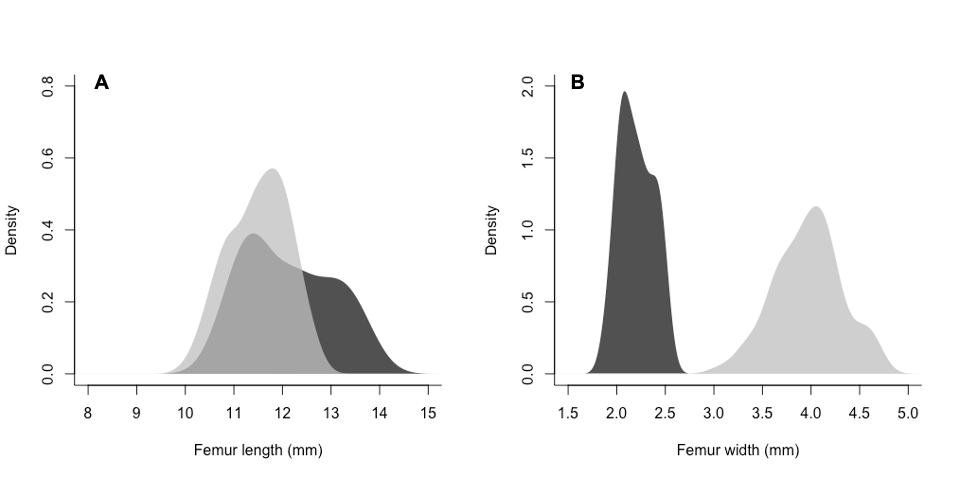

**Figure S3.** Density plots of male and female hindleg measurements. In both figures, black areas represent the female data, whereas gray areas represent the male data. **A)** Interestingly, substantial overlap in femur lengths between male and female *T. neocalinforinus*, although females in our sample had relatively longer legs. **B)** However, despite the overlap in femur length, no overall in femur width existed.

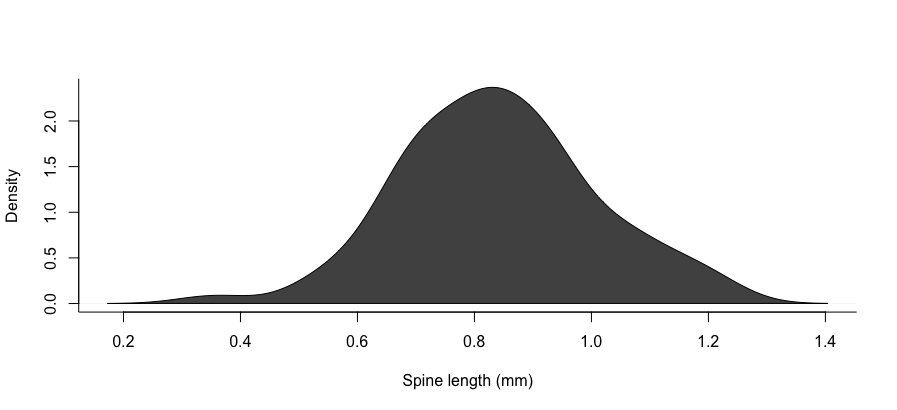

**Figure S5.** Density histogram of male prominent tibial spine length measurements

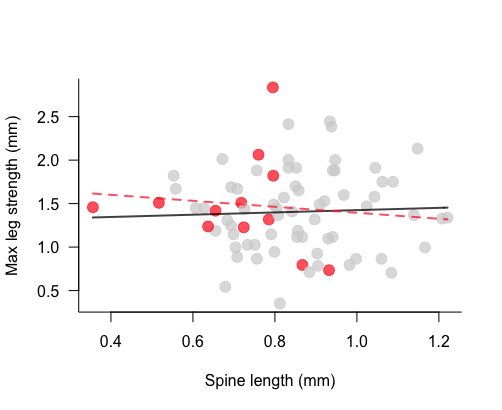

**Figure S5.**  The relationship between male tibial spine length and maximal leg force. Gray and red circles represent data from the hindlegs with a single or double tibial spine, respectively. Further, the solid black and dashed red lines represent the best fit line the relation between spine length and max leg strength for single or double tibial spine legs, respectively. In both scenarios, spine length is a poor predictor of maximal leg strength.
